# From innovation to integration: Personality, sex, and learning drive recombination of innovative behaviours in lovebirds

**DOI:** 10.64898/2026.06.25.734434

**Authors:** Shengyu Wang, Christy Yuen Ching Hung, Emily Shui Kei Poon, Simon Yung Wa Sin

**Author notes:** Correspondence or (S.Y.W.S.).

## Abstract

Behaviour innovation plays a pivotal role in a species’ adaptability to dynamic environments. Investigating innovative behaviour and its underlying mechanism is therefore crucial for elucidating the development of cognitive flexibility across animals. Animal personality—which shapes how individuals perceive and engage with their surroundings—could offer insights into individual variation in this process. This study used a three-step foraging puzzle to evaluate the innovation capacity in 28 rosy-faced lovebirds (*Agapornis roseicollis*), specifically examining their capacity to recombine individually innovated component behaviours into integrated, more sophisticated techniques. We found that nearly half of the individuals spontaneously innovated multiple component behaviours to solve novel puzzles. Crucially, when challenged with a more sophisticated task, they recombined these behaviours into functionally dependent sequences without prior social demonstration. We further identified sex, persistence, and asocial learning capacity as key predictors of innovative problem-solving performance, with females, persistent individuals, and superior asocial learners excelled at problem-solving. Our findings demonstrate that behavioural innovation is not a static event, but a dynamic process—modulated by physical, cognitive, and personality variables—in which behaviours are flexibly transferred and recombined into increasingly complex forms to enable rapid individual adaptation.

## 1. Introduction

Behavioural innovation allows animals to adapt to rapid environmental changes when genetic evolution is too slow to keep pace (Tebbich et al., 2016). Animal innovation is defined as the emergence of entirely new behaviours that provide novel solutions to environmental challenges (Reader & Laland, 2003). It is associated with adaptation to harsh environment (Roth et al., 2010), residency in urbanized landscapes (Audet et al., 2016; Sol et al., 2011), invasion success (Sol et al., 2002; Sol & Lefebvre, 2000), habitat generalism (Overington, Griffin, et al., 2011), mate preference (Chen et al., 2019), and reproductive success (Cauchard et al., 2013; Keagy et al., 2009). By conferring survival and reproductive advantages, behavioural innovations generate feedback loops in which increasingly complex behaviours selected for greater cognitive capacity. This process accelerated the pace of natural selection and speciation (e.g., Nicolakakis et al., 2003), consistent with the behavioural drive hypothesis (Wilson, 1985; Wyles et al., 1983). Thus, behavioural innovations offer a valuable source of evolutionary adaptation, and hence, a crucial mechanism for coping with environmental changes (Nicolakakis et al., 2003; Tebbich et al., 2016). Given the adaptive advantages of innovative behaviour (Lefebvre & Sol, 2008), researchers have turned to experimental approaches to uncover the proximate mechanisms driving such behaviours. By testing animals under controlled conditions, these studies have explored how cognitive and behavioural traits contribute to foraging innovations. One major emphasis of current research on animal innovation is investigating the role of random motor variability, personality differences, and learning capabilities in generating novel solutions (Griffin & Guez, 2014).

Indeed, interindividual variations in problem-solving abilities are often linked to animal personalities, the unique behaviour patterns consistent with each individual across contexts, which shape their perceptions and interactions with its environment (Dall et al., 2004; Réale & Dingemanse, 2012; Sih & Giudice, 2012). Animal personality is also termed as behavioural syndromes, which represent correlated suites of behaviours. For instance, a proactive syndrome is characterized by boldness, high exploratory tendency, and aggressiveness, whereas a reactive syndrome is defined by caution, low exploratory tendency, and shyness (Koolhaas et al., 1999; Réale et al., 2007). Such behavioural syndromes are often associated with individual traits such as sex, age, and dominance, and are further mediated by social context and life history (Stöwe et al., 2006; Wolf et al., 2007). Despite this complexity, research has revealed patterns in how personality influences problem-solving. For instance, it has been suggested that proactive individuals were linked to superior problem-solving abilities (Cauchard et al., 2013; Damerius et al., 2017; Daniels et al., 2019; Johnson-Ulrich et al., 2018; Overington, Cauchard, et al., 2011; Sol et al., 2012). However, the consistency of this relationship is questionable, and the mechanism by which personality affects problem-solving ability remains poorly understood.

Moreover, recent findings suggest that the influence of personality on cognitive performance can be contextual dependent. The speed-accuracy trade-off theory proposes that proactive individuals learn rapidly but exhibit less behavioural flexibility, whereas reactive individuals prioritize accuracy and deliberation over speed (Sih & Giudice, 2012). Empirical evidence across taxa supports this idea: for instance, fast-exploring great tits (*Parus major*) excel at initial learning but struggle with reversal tasks, while slow explorers show the opposite pattern (Verbeek et al., 1994). Similarly, aggressive jumping spiders (*Portia labiata*, Chang et al., 2018) and highly exploratory common brushtail possums (*Trichosurus vulpecula*, Wat et al., 2020) outperform in simple tasks but are often outperformed by less proactive individuals when a task becomes complex. However, the specific contextual factors that determine when a proactive or reactive personality is advantageous remain poorly understood. Factors such as the type of cognitive challenge (e.g., innovation vs. inhibition), the animal’s developmental stage, and task complexity likely influence the outcomes, though their relative contributions remain poorly understood. Furthermore, personality-performance relationships are not always consistent (Biondi et al., 2010; Bókony et al., 2014). Resolving these discrepancies requires shifting the focus from whether personality influences innovation to mapping the specific conditions—such as task type, social context, or species ecology—that determine how these relationships emerge.

Like humans, many animal species demonstrate that behavioural innovation is not static, developing progressively sophisticated behaviours by incorporating existing techniques to facilitate rapid adaptation. Originally, motor diversity, the number of distinct motor actions an animal can employ when interacting with a task or environment, provides the essential raw material for generating novel behaviours, upon which learning and experience can act to modify and refine successful variants (Griffin et al., 2014; Griffin & Guez, 2014). For example, motor diversity (e.g., pecking, pushing, grabbing, and lifting) facilitates problem-solving success and reduces solving latency in Indian mynas (*Sturnus tristis*, Griffin et al., 2014; Griffin & Diquelou, 2015). Similarly, research on spotted hyenas (*Crocuta crocuta*, Benson-Amram et al., 2013; Benson-Amram & Holekamp, 2012) and four nonhuman great ape species (Manrique et al., 2013) revealed that the flexibility of motor actions was indicative of superior innovation propensity. Grasping diverse manipulative skills should increase possibilities for learning objects’ physical properties, and provide a valuable source generating behavioural variants (Griffin & Guez, 2014). Subsequently, beyond initial acquisition, practice and experience drive the gradual perfection of behavioural techniques, leading to more proficient and efficient performance. Individual adept at learning typically enhance their performance, being more efficient and employing fewer but more effective techniques directed towards a task (e.g., Chow et al., 2016; Daniels et al., 2019). Furthermore, cognitive flexibility enables animals to access and deploy pre-existing knowledge and motor skills to navigate and solve novel challenges (Auersperg et al., 2012; Taylor et al., 2010). Finally, when confronted with complex foraging scenarios, animals can recombine elements from their behavioural repertoire to construct novel, more complex behaviours (Wild et al., 2021). However, experimental research tracing the complete pathway from innovation and individual learning to behavioural recombination in non-human animals remains scarce. Consequently, whether the ability to recombine mastered skills into a novel, functional sequence drives the emergence of complex behaviours is a critical and open question.

Parrots (Psittacidae) and corvids (Corvidae) have been termed “feathered apes” (Emery, 2004; Lambert et al., 2019) due to their exceptional cognitive capacities, which are comparable to those observed in primates (Güntürkün et al., 2017). Studies on problem-solving ability in these birds have documented exceptional performances in advanced domains such as tool manufacture (Auersperg et al., 2012), causal reasoning (Taylor et al., 2010), and cultural adaptation (Klump et al., 2021). This makes them ideal model systems for investigating the underpinnings of complex behaviour. Yet, only a limited number of Psittacidae species have been subject to comprehensive and detailed study (Auersperg & von Bayern, 2019). It remains largely unexplored whether other, especially smaller, parrot species exhibit comparable cognitive performance.

Rosy-faced lovebirds (*Agapornis roseicollis*) are highly social small-sized parrots, native to arid regions in southwestern Africa (Huynh et al., 2023), where food and water resources vary greatly among seasons (Ndithia & Perrin, 2006b). They have an apparent preference towards habitats, in which open grassland with scattered shrubs is the most preferred (Ndithia & Perrin, 2006b). They are also highly selective in diet and remote forage in groups for their frequently consumed food, *Anthephora schinzii* seeds (Ndithia & Perrin, 2006a). Species that navigate complex environmental challenges through sociality are typically expected to possess enhanced cognitive capabilities—a link proposed by several hypotheses (e.g., Social Intelligence Hypothesis, Humphrey, 1976; Jolly, 1966; Cultural Intelligence Hypothesis, Whiten & van Schaik, 2007; Cognitive Buffer Hypothesis, Allman et al., 1993; Cohen & McKay, 1984; Sol, 2008; Social Brain Hypothesis, Dunbar, 1998). Furthermore, recent evidence demonstrates that rosy-faced lovebirds exhibit sophisticated cognitive capabilities, including visual cue discrimination and inference-based numerical competence (Tsang et al., 2025; Wang et al., 2025). These birds are therefore ideal candidates for studying foraging innovation and behavioural recombination, as well as the underlying influence of physical and personality traits. Here, we utilized the rosy-faced lovebird to investigate (1) problem-solving performance using a three-step foraging puzzle; (2) inter-individual variation in four personality traits—neophobia, exploration, activity, and persistence; (3) relationship between personality traits and problem-solving performance; and (4) their ability to apply and recombine pre-existing skills to solve novel, complex tasks. This study advances our understanding of avian innovation by evaluating problem-solving performance across tasks varying in complexity, mechanical design, and motor requirements and identifying the underlying traits that govern these behaviours.

## 2. Methods

### 2.1 Study subjects and housing conditions

Twenty-eight adult (3-5 years old) rosy-faced lovebirds, consisting of 19 males and 9 females, were used for the experiment. The lovebirds were individually housed in wire-mesh cages (measuring 60 cm in length, 40 cm in width, and 40 cm in height). The room followed a light/dark cycle from 8:00 to 20:00. The ambient temperature was maintained at a constant range of 22-24°C, while the humidity remained stable at 50-60%. The lovebirds had access to artificial food pellets (Mazuri Small Bird Maintenance Diet 56A6) and UV-filtered water ad libitum.

#### Ethical note

All procedures were approved by the Committee on the Use of Live Animals in Teaching and Research (CULATR; approval number: 24-202), and under a Department of Health Animal (Control of Experiments) Ordinance Chapter 340 permit ((24-485) in DH/HT&A/8/2/3 Pt.70). A single room housed all the birds, permitting both visual and auditory contact among them. Each bird was housed singly in a cage to prevent aggression due to their territorial instincts. We promote their psychological well-being by providing environmental enrichment, such as toys, chewable items, and classical music. All lovebirds were acquired from local breeders and participated in subsequent studies after this project was completed. Their health status and mental well-being are carefully monitored daily.

### 2.2 Foraging puzzle design and innovation test

We designed three puzzles (Fig. 1a-1c), which could be combined together into a single puzzle that required birds to solve three steps successively before obtaining the food rewards (Fig. 1d). These three puzzles—lid puzzle, sliding door puzzle, and drawer puzzle—each involved distinct solving skills, movements, and solvable directions. Specifically, the lid puzzle required flipping, the sliding door puzzle needed sliding, and the drawer puzzle demanded pulling to solve. These puzzles involved up-down, left-right, and back-forward movements, respectively. Regarding the solvable directions, they could be solved through flipping at the lid’s four sides, and sliding either left or right, and unidirectional pulling only, respectively. The puzzles were made of transparent acrylic panels, which made the food reward inside visible to encourage birds to solve them.

**Figure 1.**
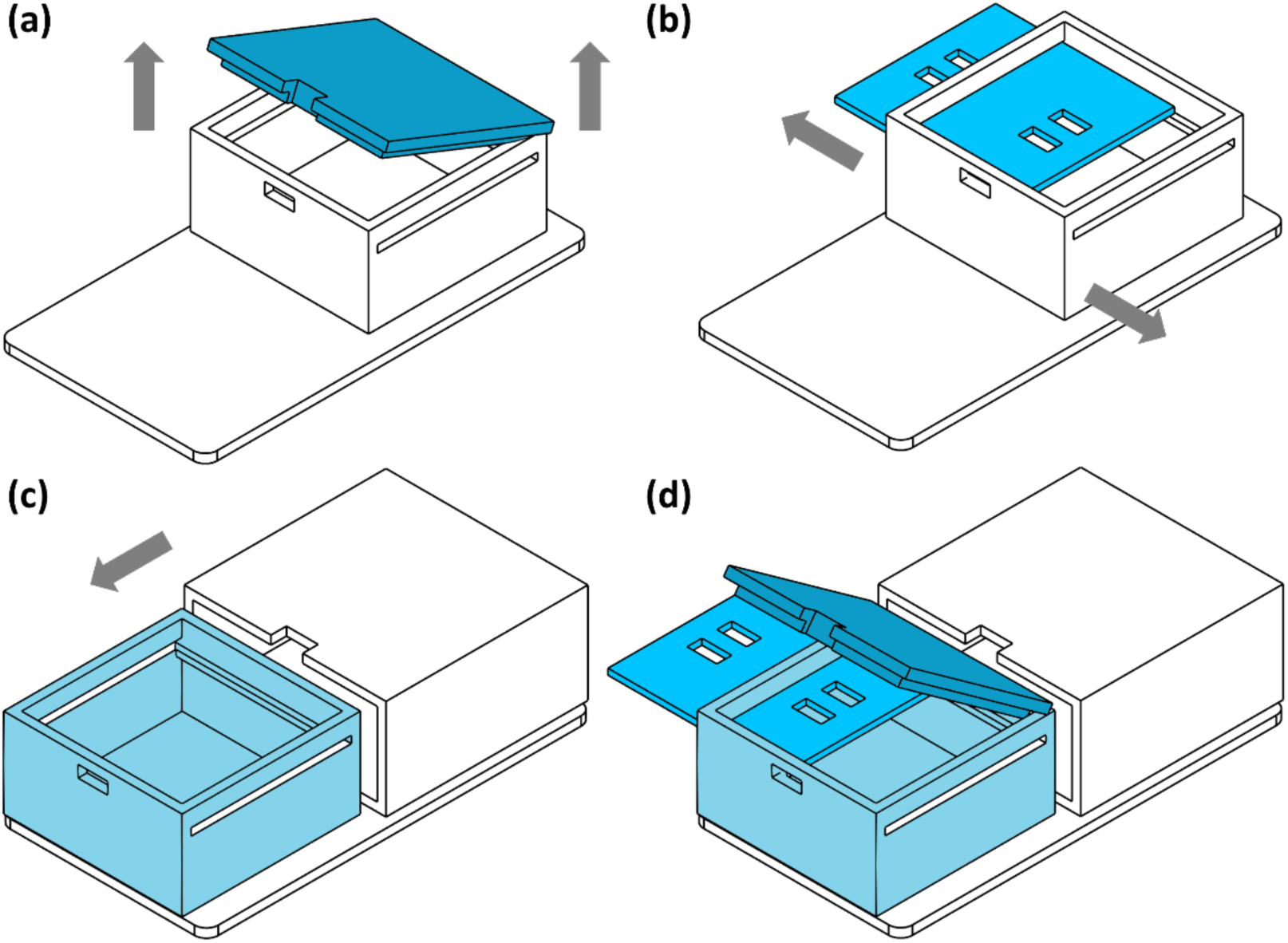
Design of the food puzzles: (a) lid puzzle, (b) sliding door puzzle, (c) drawer puzzle, and (d) combined puzzle. Puzzle dimensions: length × width × height = 6 × 6 × 3 cm. Arrows indicate the direction of puzzle-solving movements. Movable parts were coloured in each puzzle.

This study started by testing whether rosy-faced lovebirds can innovatively solve food puzzles independently on their own without prior training. All subjects were tested in their own cages, with an opaque barrier between cages during testing to prevent them seeing each other. We tested all three puzzles separately, firstly the lid puzzle, followed by the sliding door puzzle, then the drawer puzzle. Subsequently, we provided them with the combined puzzle that required solving all three steps to investigate whether they can integrate different skills and apply them to novel conditions.

To ensure all birds were motivated to solve the puzzles, we gave an acclimation period where they were allowed access to a coverless puzzle with food until all of them approached the puzzle within 30 seconds in five consecutive trials (40 acclimation trials in total). Then we conducted the innovation tests. To increase their motivation, their feeders was closed 90 minutes before and 30 minutes after the test (Wang et al., 2025). The puzzle was placed at the bottom of the cage for 30 minutes to allow them to solve it. We repeated trials until an individual a) reached the pass criterion (i.e., successfully solved a puzzle in five consecutive trials), and b) tried for at least six trials, and lost motivation (time they spent trying to solve the puzzle) by 80% compared to that in the first trial. Birds that reached the pass criterion were regarded as “solvers”, which demonstrated consistent problem-solving ability. Between the innovation test of each puzzle type, we conducted acclimation trials again as described above to ensure those birds that failed to solve the previous puzzle type regained motivation.

We conducted maximum one trial per day for each bird, alternating between morning sessions (10:00–12:00) and afternoon sessions (14:00–16:00). This practice was to avoid potential difference in motivation at different times of a day. The foraging puzzle contained a mixture of seeds and food pellets as food rewards. Cameras (Mi Camera 2K, Xiaomi Communications Co., Ltd.) were placed on the top of the cage to record the behaviour of the birds for analysis.

### 2.3 Personality test

We measured four aspects of personality—neophobia, exploration, activity, and persistence—to investigate the correlation between innovativeness and personality. Each trial of neophobia, exploration, and persistence tests applied the same practice of 90-minute pre-trial fasting, 30-minute test trial, and 30-minute post-trial fasting and the same food rewards as in the innovation tests.

#### 2.3.1 Neophobia

Neophobia is defined as the reluctance or aversion to approach novel objects (Greenberg, 2003). To assess this behaviour, we measured the latency to feed near novel objects compared to the baseline feeding latency (i.e., without novel stimuli) (Greenberg & Mettke-hofmann, 2001). Specifically, neophobia was quantified as the difference between the mean latency to feed from a familiar feeder (i.e. a puzzle without cover) placed beside a novel object (Fig. 2b-2d) across three trials and the mean baseline latency (Fig. 2a) from three trials without novel objects. In the case when an individual did not approach the food rewards within a trial, the maximum latency (i.e. 30 minutes) was recorded. To determine the consistency of neophobic response across different stimuli, we used three novel objects (Fig. 2b-2d) varying in colours, textures, and shapes. The neophobia score for each individual was calculated as the average of their neophobic responses (i.e., the latency differences) across all three novel objects.

**Figure 2.**
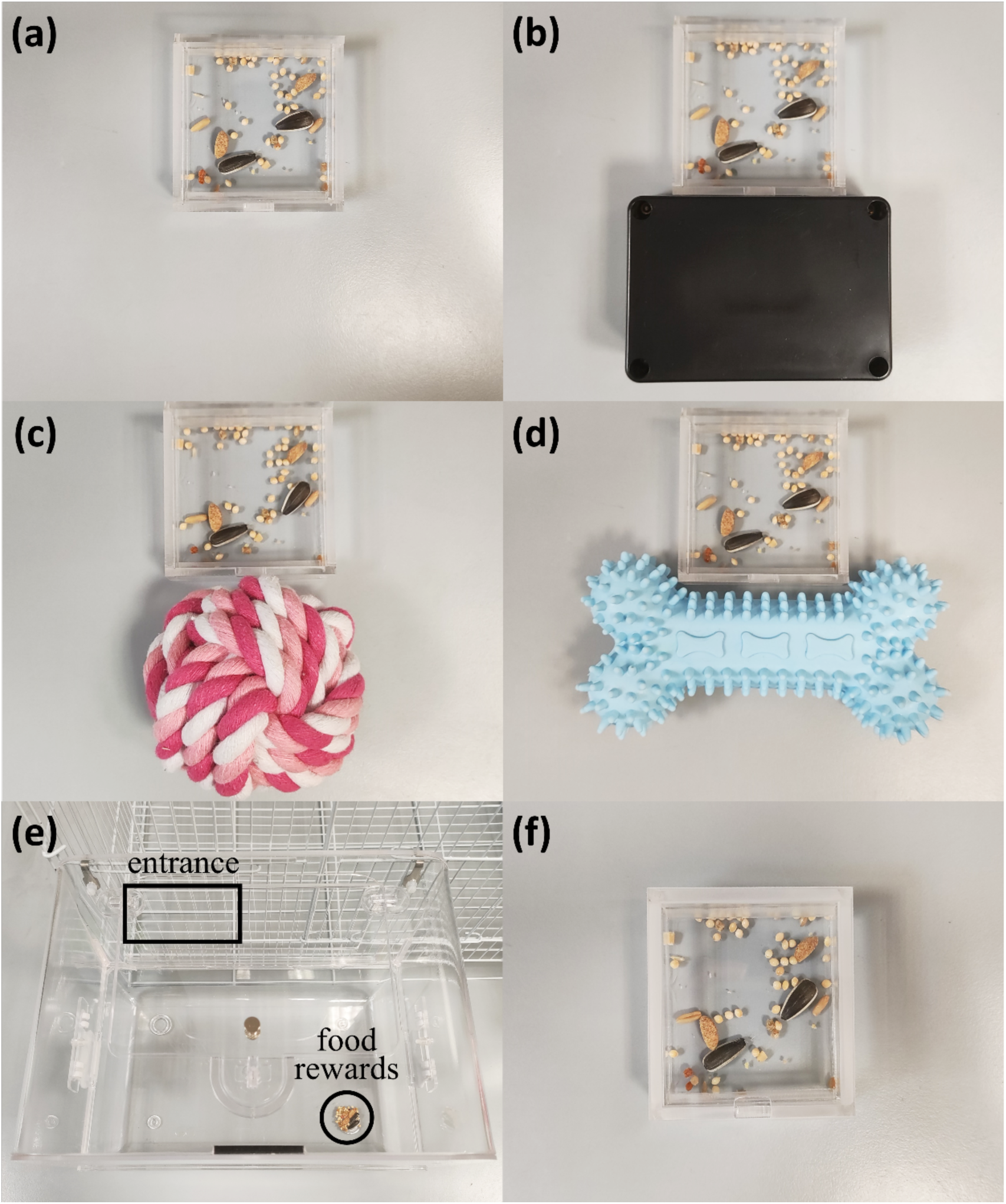
Experimental setup for personality tests. a) Neophobia control test: open puzzle containing free access food rewards. b) Neophobia test object 1: black plastic box (length × width × height = 8.3 × 5.8 × 3.4 cm). c) Neophobia test object 2: pink-white cotton knot ball (diameter: 7 cm). d) Neophobia test object 3: blue rubber bone (length × width × height = 12 × 6 × 3.5 cm). e) Exploration test: external chamber attached to the cage, with food rewards (circled) at the opposite corner of the entrance (squared). f) Persistence test: sealed puzzle.

#### 2.3.2 Exploration

We conducted the exploration test by attaching an external chamber (length × width × height = 30 × 21.5 × 22.5 cm) to their cages with food rewards placed at the far corner opposite the entrance (Fig. 2e). To obtain the food, the lovebirds had to enter the chamber. Birds that did not access the food rewards during the trial had latency recorded as 30 minutes. The exploration score was calculated by subtracting the mean latency to feed from the testing time (i.e. 30 minutes) across three trials.

#### 2.3.3 Activity

Activity was assessed by measuring the mean frequency of movement during a 10-minute observation window recorded by a camera set at the front of the cage. Three trials were conducted, all between 10:00–11:00 to control for potential diurnal variations in activity levels. The activity score was defined as the average movement frequency across the three trials.

#### 2.3.4 Persistence

We evaluated persistence by the subjects’ continuity of attempting action towards the foraging puzzle (Griffin & Guez, 2014). We covered the puzzle with a transparent lid that could not be opened (Fig. 2f) and the lovebirds were given 30 minutes per trial (three trials in total) to open it. The time spent trying to access the food was recorded. The persistence score was calculated as the mean attempting time across the three trials.

### 2.4 Data collection

We collected the behavioural data using the software Behavioral Observation Research Interactive Software (BORIS) v.8.25 (Friard et al., 2016) during video analysis. The data to collect were latency to the first attempt (i.e., the time in seconds the subjects entered the puzzle region for the first time), latency to solve (i.e., the time in seconds they ate the first food reward), solving time (i.e., latency to solve minus latency to the first attempt), attempting time (i.e., the total duration in seconds spent by the subjects within the confines of the puzzle region), success rate (i.e., the number of successful trials divided by the total number of trials for an individual within each puzzle type), and success iterations (i.e., the ordinal number of successful trials; for example, a success iteration of 1 indicates the first success, regardless of previous attempts). For the neophobia and exploration tests, we measured latency to feed (i.e., the time in seconds until the first food reward was eaten). For the activity test, we recorded the proportion of time that they moved actively. For the persistence test, we measured the attempting time.

As a measure of asocial learning ability, we used a learning score calculated as the mean solving time of each bird across the four trials after their first success (Overington, Cauchard, et al., 2011). This calculation reflects the birds’ improvement in puzzle-solving proficiency following their initial success. The slope of the learning curve was not employed due to several fast solvers immediately solved the puzzle in the following trial after the first success, resulting in a curve with a slope of zero. For birds that did not solve the puzzles, the learning score was recorded as NA.

### 2.5 Statistical analysis

We used RStudio (Version 2024.09.0+375) (R Core Team, 2022) for statistical analysis and figure plotting. Evaluation of problem-solving capabilities was based on the puzzle-solving outcome (success/failure) of each individual and the solving time of successful individuals. To compare success rates and solving times across different puzzles, we first conducted a Kruskal-Wallis test. Subsequently, we performed post hoc pairwise comparisons (Dinno, 2024) to identify specific differences. The Wilcoxon rank sum test was used for comparing the solving time between different success trials.

We used Pearson correlation tests to evaluate the correlation coefficients and their corresponding significance among personality traits (neophobia, exploration, activity, and persistence) and life-history factors (sex and age). Independent samples t-tests were conducted to examine sex differences for each of the four personality traits.

We used generalised linear mixed models (GLMMs) (Bates et al., 2015) to investigate the association between problem-solving performance and personality and sex, with puzzle type (i.e., lid puzzle, sliding door puzzle, drawer puzzle, and combined puzzle) and bird identity (ID) as random effects. Age was excluded from the analysis due to limited variance, since all birds were of similar ages. Besides, activity was not included in the analysis due to its strong correlation with sex (Fig. S1). The following three models were constructed. Model 1 is a GLMM with Bernoulli distribution testing the association between variables and subjects’ success: Success ∼ sex + trial + neophobia + exploration + persistence + (1| puzzle type) + (1|ID). Model 2 is a GLMM with gamma distribution testing how sex, behavioural traits and task-specific experience predict problem-solving efficiency in successful attempts: Solving time ∼ sex + success iterations + success rate + neophobia + exploration + persistence + (1|puzzle type) + (1|ID). Solving time was recorded as NA when the subject failed in an experiment, and thus was not included in Model 2. On the basis of Model 1, we further included learning score as one variable to evaluate whether asocial learning ability could predict success among those who made at least one success, i.e., Model 3: Success ∼ sex + trial + neophobia + exploration + persistence + learning + (1| puzzle type) + (1|ID). Neophobia score, exploration score, activity score, persistence score, and learning score were standardized by subtracting the mean and then dividing by the standard deviation. All assumptions of the statistical models were assessed and met.

## 3. Results

### 3.1 Problem-solving performance across different puzzle types

Twenty-eight lovebirds participated in solving the four puzzles: 13 (46.43%), 22 (78.57%), 19 (67.86%), and 11 (39.29%) birds successfully solved the lid puzzle, the sliding door puzzle, the drawer puzzle, and the combined puzzle, respectively (Table 1; Video S1). Five males were unable to solve any puzzles, while all other individuals successfully solved at least one puzzle. Twelve individuals successfully solved all three component puzzles, and nine of them also solved the combined puzzle (Fig. 3). We conducted 11, 17, 13, and 7 trials for the lid, sliding door, drawer, and combined puzzles, respectively, continuing each set until the individual either reached the pass criterion or lost motivation (Table 1). The lovebirds required the fewest trials to achieve their first success on the combined puzzle, followed by the drawer, sliding door, and lid puzzles (Table 1). A similar pattern was observed for reaching the pass criterion, with the combined puzzle again requiring the fewest trials, followed by the same order of puzzles (Table 1).

**Figure 3.**
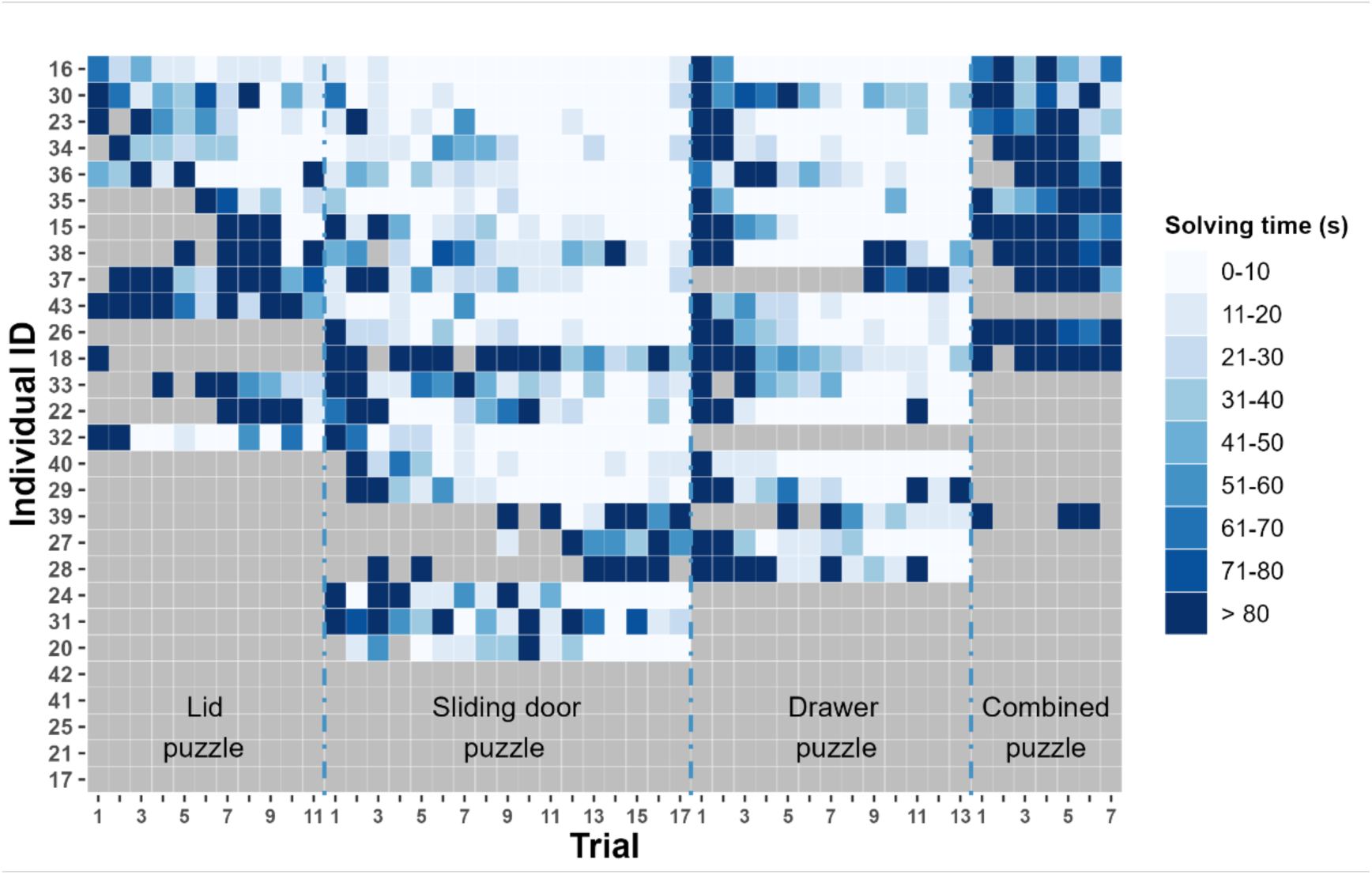
Performance of each individual in different trials using the lid, sliding door, drawer, and combined puzzles. Individuals are ranked by their average success rate across four puzzle types, with the most successful individual at the top. When success rates are equal, they are ranked by shorter to longer mean solving time from the top to bottom. The intensity of blue indicates their solving time in each trial. Grey box indicates failure in solving the puzzle.

**Table 1.** Puzzle-solving performance for the four puzzles, indicated by the total trials, number of solvers, mean first success, mean trials to pass, mean success rate, and mean solving time.

|  | Total<br>trials | Number of<br>solvers (%) | Mean first success<br>± SEM # | Mean trials to pass<br>± SEM * | Mean success rate (%)<br>± SEM | Mean solving time (s)<br>± SEM # |
| --- | --- | --- | --- | --- | --- | --- |
| Lid puzzle | 11 | 13 (46.43%) | 2.86 ± 0.64 | 7.46 ± 0.74 | 37.34 ± 8.13 | 228.33 ± 52.21 |
| Sliding door puzzle | 17 | 22 (78.57%) | 1.91 ± 0.48 | 6.68 ± 0.71 | 74.37 ± 7.49 | 82.24 ± 32.2 |
| Drawer puzzle | 13 | 19 (67.86%) | 1.63 ± 0.46 | 5.84 ± 0.51 | 64.01 ± 8.84 | 90.31 ± 20.16 |
| Combined puzzle | 7 | 11 (39.29%) | 1.50 ± 0.23 | 5.73 ± 0.27 | 37.24 ± 8.55 | 280.56 ± 65.22 |

SEM: standard error of the mean.
\* Birds that did not meet the pass criterion were excluded from the calculation of mean trials to pass.

Four puzzles varied in both success rate (Fig. 4a) and solving time (Fig. 4b). The successful rates of the lid and combined puzzles were significantly lower (*p* < 0.01) than those of the sliding door and drawer puzzles (Fig. 4a). The solving time of the lid and combined puzzles were significantly longer (*p* < 0.05) than that of the sliding door and drawer puzzles (Fig. 4b). Overall, puzzles with lower success rates required more time to solve: the combined puzzle exhibited the lowest success rate and the longest mean solving time, followed by the lid, the drawer, and finally the sliding door puzzles (Table 1).

**Figure 4.**
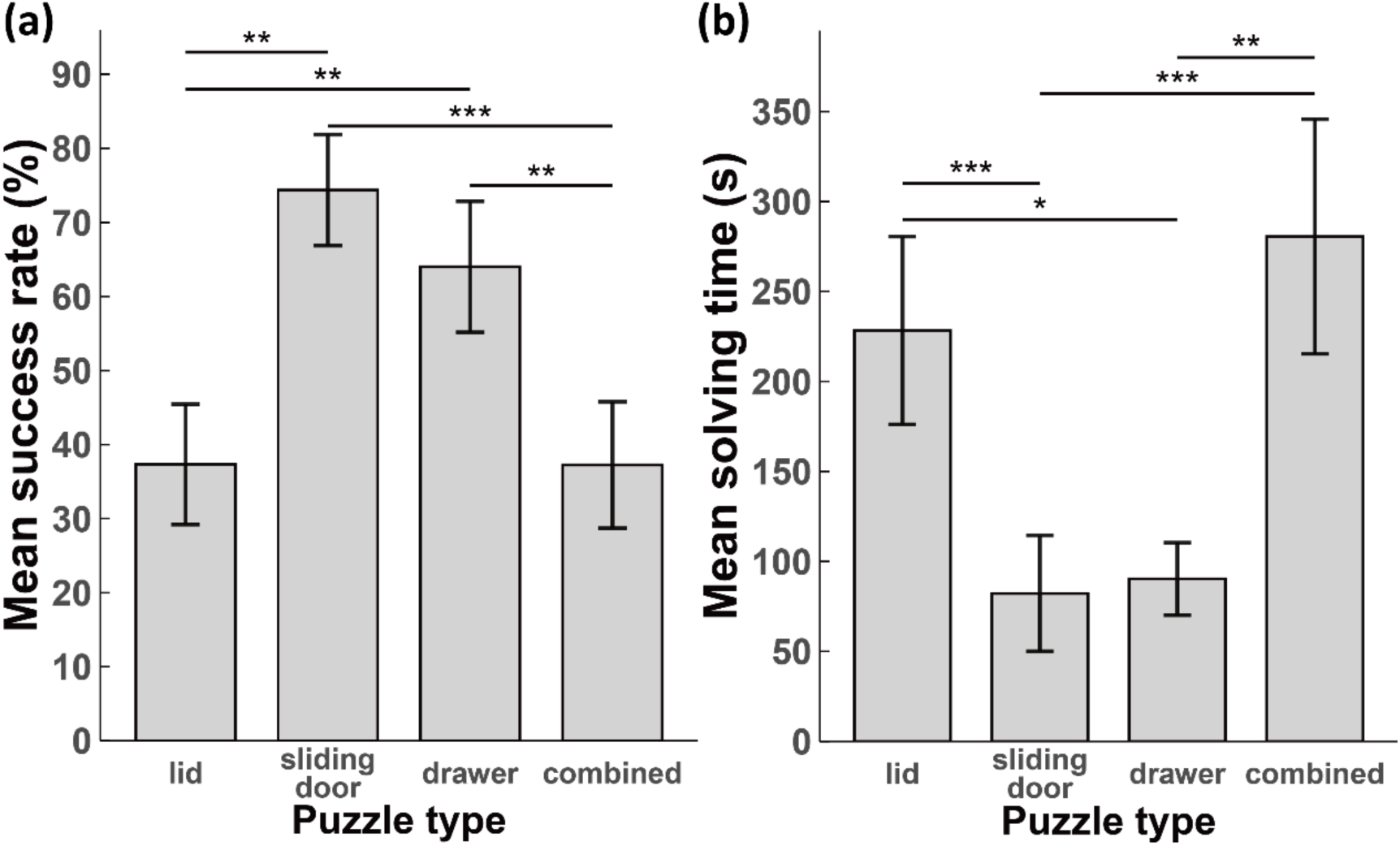
The (a) mean success rate and (b) mean solving time for each puzzle. Bars represent the mean values, and error bars indicate the standard error of the mean (SEM). SEM: standard error of the mean. * *p* < 0.05; ** *p* < 0.01; *** *p* < 0.001.

### 3.2 Correlations and sex differences in multifaceted personality traits

Highly positive correlations were evident across the three neophobia tests (object 1 and 2: *r* = 0.616; object 1 and 3: *r* = 0.720; object 2 and 3: *r* = 0.636; all with *p* < 0.001), suggesting a consistent tendency of aversion across different novel objects in an individual. Significant negative correlations were found between the neophobia score and exploration score (*r* = -0.396, *p* = 0.037), and between neophobia score and activity score (*r* = -0.419, *p* = 0.026), suggesting neophobic individuals tended to be less exploratory and less active. Persistence did not correlate with any other personality traits. The t-test result revealed a significant activity score difference between female and male [t(17.535) = 3.328, *p* = 0.004; Fig. S1], where females (mean = 0.340, SD = 0.091) demonstrated significantly higher activity levels compared to males (mean = 0.212, SD = 0.102). No significant sex differences were found for the other three personality traits (*p* > 0.05).

### 3.3 Key drivers of puzzle-solving success: sex, trial number, and persistence

Model 1, which tested the influential factors on puzzle-solving success, revealed that sex (*β* = -2.298, *p* = 0.037), trial number (*β* = 0.179, *p* < 0.001), and persistence (*β* = 3.206, *p* < 0.001) significantly influenced the success of puzzle-solving (Table 2). Specifically, females demonstrated a significantly higher success rate than males (Fig. 5a), outperforming them across all four types of puzzles (Fig. S2a). The likelihood of success increased with the number of trials (Fig. 3) and persistence (Fig. 5b). Less neophobic (*β* = -0.579, *p* = 0.266; Fig. S3a) and more explorative (*β* = 0.130, *p* = 0.810; Fig. S3b) individuals had a higher tendency of success, but the effects were not significant.

**Figure 5.**
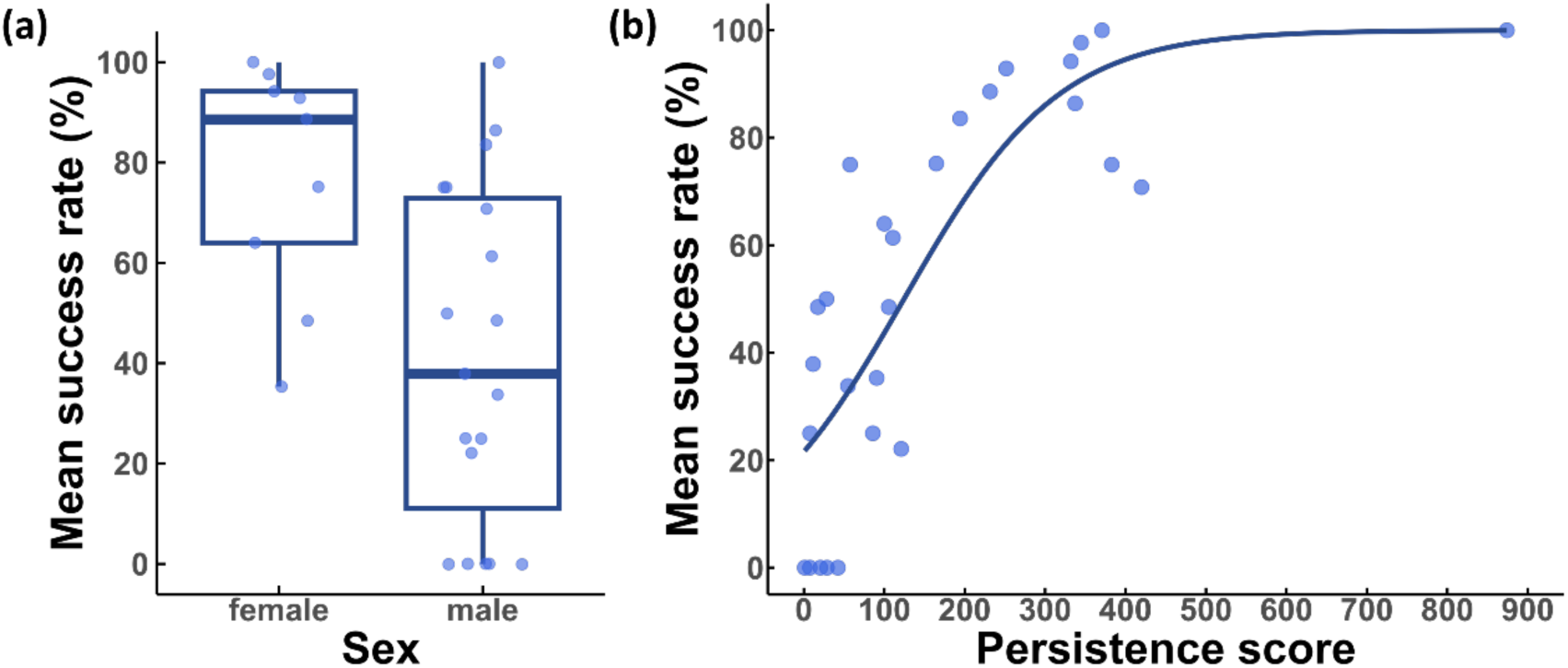
The associations of the mean puzzle-solving success rate of each individual across four puzzles with (a) sex, and (b) persistence score.

**Table 2.** Summary of GLMMs results investigating association between factors and puzzle-solving outcome (success/failure) (Model 1 & Model 3) or solving time (Model 2).

| GLMMs | explanatory variables | $\beta$ | SE | z | p | SD of random effects | |
| --- | --- | --- | --- | --- | --- | --- | --- |
|  |  |  |  |  |  | puzzle type | Bird ID |
| Model 1 | sex (male) | -2.298 | 1.099 | -2.091 | **0.037** | 1.563 | 2.319 |
|  | trial number | 0.179 | 0.027 | 6.527 | **<0.001** |  |  |
|  | neophobia | -0.579 | 0.520 | -1.113 | 0.266 |  |  |
|  | exploration | 0.130 | 0.539 | 0.240 | 0.810 |  |  |
|  | persistence | 3.206 | 0.787 | 4.076 | **<0.001** |  |  |
| Model 2 | sex (male) | 0.565 | 0.246 | 2.295 | **0.022** | 0.817 | 0.819 |
|  | success iterations | -0.183 | 0.009 | -20.760 | **<0.001** |  |  |
|  | success rate | -1.192 | 0.321 | -3.714 | **<0.001** |  |  |
|  | neophobia | 0.130 | 0.129 | 1.002 | 0.316 |  |  |
|  | exploration | -0.146 | 0.136 | -1.077 | 0.282 |  |  |
|  | persistence | -0.202 | 0.125 | -1.614 | 0.107 |  |  |
| Model 3 | sex (male) | 0.574 | 0.653 | 0.879 | 0.379 | 0.774 | 1.153 |
|  | trial number | 0.387 | 0.052 | 7.388 | **<0.001** |  |  |
|  | neophobia | 0.067 | 0.353 | 0.191 | 0.848 |  |  |
|  | exploration | 0.358 | 0.386 | 0.928 | 0.353 |  |  |
|  | persistence | 0.603 | 0.434 | 1.391 | 0.164 |  |  |
|  | learning | 1.420 | 0.216 | 6.569 | **<0.001** |  |  |

Significant *p* values are in bold.

### 3.4 Determinants of solving efficiency: sex, success iterations, and success rate

Model 2 that examined factors associating with solving time indicated that sex, success iterations, and success rate influenced puzzle-solving time (Table 2). Specifically, females solved the puzzles significantly faster than males (*β* = 0.565, *p* = 0.022; Fig. 6a; Fig. S4), achieving a shorter average solving time across all four puzzles (Fig. S2b). Solving time decreased with an increase in the number of successful attempts (*β* = - 0.183, *p* < 0.001; Fig. 6b; Fig. S4) and a higher success rate (*β* = -1.192, *p* < 0.001; Fig. 6c). No significant effects were detected for any personality traits on solving time.

**Figure 6.**
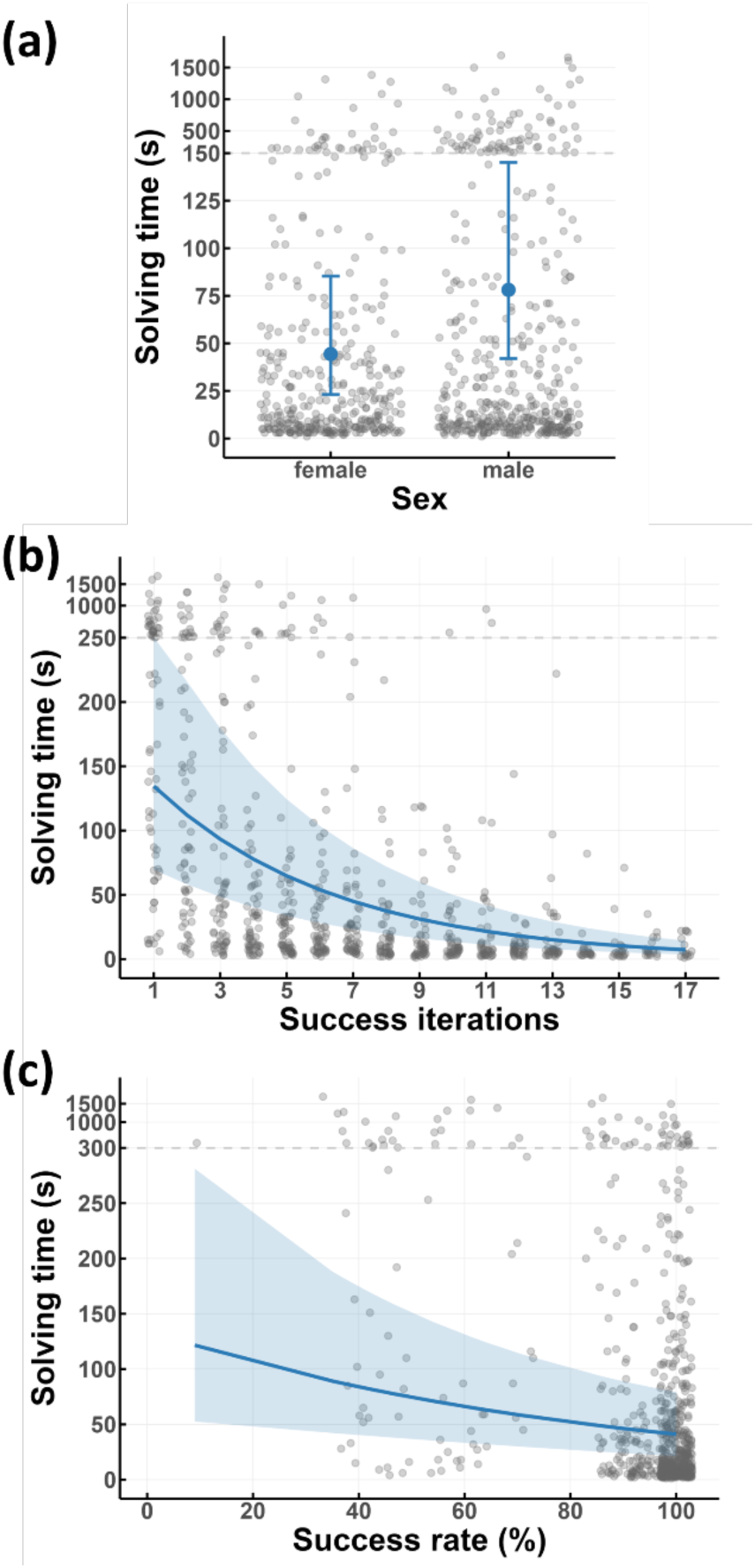
The associations between puzzle-solving time and (a) sex, (b) success iterations, and (c) success rate. To accommodate the full data range, the y-axis scale is compressed 30-fold for values exceeding 150 (a), 250 (b), and 300 (c).

### 3.5 Practice makes proficient: asocial learning and performance improvement

As the cumulative count of successful attempts increased, individuals improved in their puzzle-solving performance. Compared with the first success, a substantial decrease in average solving time was observed in the 4th success for the lid puzzle (dropped by 77.66%), the 4th success for the sliding door puzzle (dropped by 84.51%), 2nd success for the drawer puzzle (dropped by 67.05%), and the 6th success for the combined puzzle (dropped by 55.48%; Fig. S4a). Model 3 further revealed that lovebirds with better asocial learning ability (i.e., higher learning score) had higher success likelihood (*β* = 1.420, *p* < 0.001; Fig. S3c).

## 4. Discussion

Our study revealed remarkable innovative problem-solving abilities in rosy-faced lovebirds, with notable individual variation: females and individuals characterized by high persistence and faster asocial learning performed better across different tasks. Most interestingly, we demonstrated that lovebirds are capable of skill cumulation—spontaneously recombining innovative component behaviours into complex, integrated sequences. This study provides systematic insights into the origins and evolution of complex animal behaviour, as well as the factors contributing to such processes.

Lovebirds in this study demonstrated remarkable and rapid problem-solving abilities, a proficiency that not only rivals the most intelligent avian groups but also suggests that innovative capacity is a conserved trait within parrots, strongly linked to relative brain size (Gutiérrez-Ibáñez et al., 2018). Regardless of puzzle complexity, a large proportion of lovebirds successfully solved the tasks. Even more impressively, they typically achieved their first success within the first three trials, and the vast majority continued to succeed thereafter. The high problem-solving proficiency observed in this study firmly positions lovebirds among the most adept avian innovators, comparable to those of corvids and large parrots, and potentially superior to other avian species and non-primate mammals. For instance, in Goffin’s cockatoos (*Cacatua goffiniana*), the proportions of successes for the slide task and cover task were approximately 60%, while that for the drawer task was around 80% (Rössler et al., 2020). Similarly, 5 out of 6 ravens (*Corvus corax*) solved vertical pull-up task (Heinrich & Bugnyar, 2005). This suggests that the capacity for innovation propensity is likely highly conserved throughout the parrot family, irrespective of body size or absolute brain volume. Furthermore, the demonstrated innovation capacity in Psittacidae and Corvidae appears to be markedly superior to that observed in other bird families. Great tits solved lever-pulling tasks at rates of 44% (overnight) and 26% (daytime), while 25% mastered daytime string-pulling (Cole et al., 2011). Similarly, 55.56% of Carib grackles (*Quiscalus lugubris*) opened a hinged box within eight trials (Overington, Cauchard, et al., 2011), and urban bullfinches (*Loxigilla barbadensis*) outperformed rural ones (50% vs. 26%) on a tunnel task (Audet et al., 2016). Similar patterns emerge in non-primate mammals: 20.63% of meerkats (*Suricata suricatta*) solved at least one of three lid tasks, though only 6.35% solved all three (Thornton & Samson, 2012), while 14.5% of hyenas successfully opened a latch bolt (Benson-Amram & Holekamp, 2012). Cross-species comparative analyses indicate that relative brain volume may underpin sophisticated cognitive processes (Cnotka et al., 2008; Mehlhorn et al., 2010) and innovative behaviours (Lefebvre et al., 2004).

Substantial research indicates that interspecific variation in innovation reflects differences in cognitive capacity. In lovebirds, superior problem-solving capacity is likely an evolutionary adaptation shaped by their complex social and foraging ecology. Surviving in semi-arid environments with sparse, scattered, and highly seasonal food resources (Ndithia & Perrin, 2006a, 2006b), the ability to flexibly solve problems—such as manipulating novel food sources or navigating a changing landscape—is critical. This pressure to locate ephemeral resources is consistent with established ecological patterns, which suggest that species in volatile niches tend to develop more flexible behavioural strategies (Davey, 1989; Day et al., 1999). Furthermore, their intricate social structure—characterized by social foraging and colonial nesting (Ndithia et al., 2007; Ndithia & Perrin, 2006a)—intensifies cognitive demands. In this competitive setting, individuals who can innovate new foraging techniques, defend resources efficiently, and learn from others would gain a significant survival advantage. This aligns with the Social Intelligence Hypothesis (Humphrey, 1976; Jolly, 1966) and the related Social Brain Hypothesis (Dunbar, 1998), which posit that complex social life is a primary driver of cognitive evolution.

On the individual scale, lovebirds with advanced problem-solving abilities consistently outperform their counterparts. This superiority is demonstrated through three key findings: individuals who excel at one task also tend to perform well on others (Fig. 3), showing a consistent problem-solving ability across domains; those who are more likely to solve a task also do so more quickly (Model 2); and birds that refine their techniques faster have a higher probability of success (Model 3). Collectively, these results point to the existence of enduring individual differences in innovative propensity. A meta-analysis highlights widespread, consistent individual differences in cognitive performance across taxa, suggesting a link to fitness outcomes (Cauchoix et al., 2018). This discovery parallels observations in the guppy, *Poecilia reticulata*, where past innovators displayed a higher propensity for innovation compared to past non-innovators (Laland & Reader, 1999). One plausible explanation is that specific intrinsic individual traits may predispose certain individuals to be notably inclined towards innovation. In great tits, both field and laboratory studies have demonstrated enduring stability in individual propensities to tackle unfamiliar food extraction challenges (Cole et al., 2011; Morand-Ferron et al., 2011). These enduring propensities towards innovation may interrelate with other behavioural attributes, potentially constituting a component of a behavioural syndrome (Sih et al., 2004) or personality (Dall et al., 2004).

Lovebirds showed a non-significant trend where individuals with lower neophobia and higher exploration levels were more likely to solve the problem. This trend is aligned with a body of literature linking boldness and reduced fear of novelty to enhanced innovative problem-solving in species like great tits (Cauchard et al., 2013), grackles (Overington, Cauchard, et al., 2011), pigeons (*Columba livia*, Bouchard et al., 2007), and house sparrows (*Passer domesticus*, Bókony et al., 2014). However, the fact that this relationship was not statistically significant in our experiment provides a crucial insight. Novelty responses do not consistently correlate with problem-solving (e.g., Aplin et al., 2013; Boogert et al., 2008; Morand-Ferron et al., 2011), it could represent a contextual factor influencing how animals engage with novel situations, thereby impacting problem-solving opportunities (Griffin & Guez, 2014; Reader & Laland, 2003). The non-significance of neophobia and exploration in our study possibly due to lovebirds becoming habituated to the apparatus after acclimation, leading birds of various neophobic and exploration levels to be similarly willing to engage in the task. This explanation is supported by research on Indian mynas demonstrating that the predictive power of neophobia on problem-solving can diminish as animals become familiar with a task (Griffin et al., 2013, 2014; Sol et al., 2012). Thus, the influence of novelty responses on problem-solving may be due to a shared underlying mechanism: individuals that prefer familiar and secure environments tend to exhibit a decreased propensity for feeding motivation or enhanced stress (Griffin & Guez, 2014; Sol et al., 2012; Webster & Lefebvre, 2001).

Divergent from the contextual effect of novelty responses on problem-solving, our result underscores the pivotal significance of persistence in problem-solving proficiency, where individuals that engage more with the task are more inclined to solve. This trend has been observed in various species such as meerkats (Thornton & Samson, 2012), spotted hyenas (Benson-Amram & Holekamp, 2012), woodpecker finches (*Cactospiza pallida*, Tebbich et al., 2010), Indian mynas (Griffin et al., 2014; Sol et al., 2012), great tits (Cauchard et al., 2013; Morand-Ferron et al., 2011), and blue tits (*Cyanistes caeruleus*, Morand-Ferron et al., 2011), where the likelihood of problem-solving success escalates with prolonged interactions with novel apparatuses. These findings underscore the critical role of task-oriented motivation in the problem-solving process, as individuals with greater intrinsic interest in the task tend to engage more actively. This heightened involvement potentially facilitates a deeper understanding of the task’s properties and underlying mechanisms, thereby increasing the likelihood of successful outcomes. This suggests a role of necessity in facilitating problem-solving. The “necessity drives innovation hypothesis” (Reader & Laland, 2001) posits that individuals are more inclined to invest in potentially costly innovative and social learning endeavours when dissatisfied with their current circumstances. While previous research has identified food deprivation as a key necessity enhancing persistence (Griffin et al., 2014), our methodology, which measured persistence as continued effort under equivalent conditions, indicates that intrinsic personality differences are equally pivotal. The variation in persistence scores among our lovebirds, independent of immediate hunger, suggests that individuals possess baseline differences in how they respond to frustration and challenge when facing novel tasks.

Despite that heightened persistence promotes problem-solving success in lovebirds, its impact on problem-solving efficiency is only close to marginal (Model 2). One plausible explanation is that after initially solving the puzzle, continued practices allows asocial learning to become the key factor driving their problem-solving performance. Our finding shows that individuals who improved faster after the initial success (as captured by the learning score in Model 3) had higher possibility of success.

This aligns with findings in other avian species demonstrating direct association between proficiency in extractive foraging tasks and the process of learning (Boogert et al., 2008; Bouchard et al., 2007; Overington, Cauchard, et al., 2011). Problem-solving performance consistently links to the speed of learning, when learning involves the gradual acquisition of motor techniques, which subsequently determines the repeatability and proficiency of innovation. This observation implies a connection between problem-solving abilities and operant learning, a process in which behaviours leading to desired outcomes are reinforced while those that do not are discouraged (Thorndike, 2013). Upon the generation of a motor variant, operant learning guarantees the reinforcement of effective variants; such experience further shapes the expression of motor skills, and optimizes subsequent problem-solving endeavours (Bayern et al., 2009; Griffin & Guez, 2014; Manrique et al., 2013; Reader & Laland, 2003).

Another primary discovery in this study revolves around the disparities in problem-solving capabilities between sexes, with females consistently outperforming males across three key dimensions. First, females achieved higher success rates than males in all four puzzles (Fig. S2a). Second, females completed the puzzles more quickly than males across all tasks (Fig. S2b). Third, females exhibited more consistent improvement in performance with increasing success (Fig. S4b); while males also enhanced their solving efficiency through repeated success, their solving times showed considerably greater fluctuation (Fig. S4c). Previous research has documented sexual variations in problem-solving ability across various species. However, the advantage of innovation does not consistently favour one sex across taxonomic groups. For instance, male meerkats demonstrate superior problem-solving abilities compared to their female counterparts (Doolan & Macdonald, 1996; Thornton & Samson, 2012). Across all primates, there exists a higher occurrence of innovation among males than females (Reader & Laland, 2001). In the guppy, females demonstrated a higher likelihood of innovation compared to males, a trend that could be attributed to asymmetries in parental investment between the two sexes (Laland & Reader, 1999). Juvenile female blue tits demonstrated a twofold higher likelihood of acquiring the novel skill compared to other avian counterparts, possibly due to female blue tits’ elevated nutritional requirements for heightened reproductive investment (Aplin et al., 2013). Thus, sex-based differences in problem-solving may have evolved as adaptations to the distinct survival and reproductive demands faced by males and females. Proximate mechanisms, such as hormonal activation (e.g., Galea et al., 1996), early development (e.g., Williams & Meck, 1991), and neuroanatomical substrates (Juraska, 1991), may also contribute to sexual differences in cognitive performance.

In our case, it is plausible that the enhanced problem-solving abilities observed in females are attributable to the cognitive demands associated with their distinct reproductive functions. As the primary nest builders, females rosy-faced lovebirds are responsible for the complex task of sourcing, processing, and transporting specific materials like bark strips, twigs, and thorns from identified tree species to construct a secure, cup-shaped nest within a cavity (Eberhard, 1998; Ndithia et al., 2007). This intricate process requires sophisticated spatial reasoning, material manipulation, and trial-and-error learning. This pronounced sex-specific role was further evidenced in our study by a marked difference in activity level, with females demonstrating significantly greater physical movement while scouting for nest material, building the nest, eating, and drinking. All of these contribute to a general problem-solving capacity, which is itself further refined and enhanced through accumulated experience and growing proficiency (Reader & Laland, 2001). This sex-specific behavioural ecology suggests that superior problem-solving in female lovebirds is an adaptive trait directly linked to reproductive success. Overall, our results supports the hypothesis that sex differences in innovation vary across species depending on the resource-seeking motivations of each sex (Reader & Laland, 2001).

Our study provides the first systematic demonstration of an avian species innovating skills to solve a novel challenge and recombining these acquired techniques into functionally dependent sequences to solve a sophisticated task. By showing that a single bird could independently acquire, chain, and execute distinct actions into a compound solution, this research offers robust empirical evidence for cumulative complex behaviour. The findings reveal that this capacity is not uniquely human (Dean et al., 2012). Instead, it can emerge through purely individual innovation, requiring neither social learning (Mesoudi & Thornton, 2018) nor external scaffolding and progressive rewards (Wild et al., 2021). Our findings imply that the evolutionary roots of cumulative culture lie in individual innovation and cognitive recombination, with social learning serving to amplify and accelerate these pre-existing individual capacities (Reindl et al., 2020).

## Conclusion

Rosy-faced lovebirds demonstrated remarkable innovative problem-solving abilities, comparable to those observed in corvids and large parrots, suggesting convergent evolution of this cognitive capacity in certain taxonomic groups. We identified the key traits that underpin this innovative capacity: problem-solving success is associated with sex, persistence, and asocial learning capacity, highlighting a core suite of traits that enable individuals to innovate and recombine novel behaviours. Crucially, our study provides a framework for understanding the emergence of complex behaviours. The findings establish that innovation and the recombination of individually acquired component behaviours represent a route to behavioural flexibility and complexity. This capacity may not only enable rapid individual adaptation but also serve as precious raw material in the population’s skill repository, supporting long-term behavioural evolution.

## Author Contributions

**Shengyu Wang:** Writing — review & editing, Writing — original draft, Visualization, Methodology, Investigation, Formal analysis. **Christy Yuen Ching Hung:** Investigation. **Emily Shui Kei Poon:** Writing — review & editing, Resources. **Simon Yung Wa Sin:** Writing — review & editing, Supervision, Resources, Project administration, Methodology, Investigation, Funding acquisition, Conceptualization.

## Data Availability

Data and code are available online as supplementary materials.

## Acknowledgements

We would like to express our gratitude to CCMR staff Anson Cheuk Ming Cheung, Mei Ying Wu, Shuk Kuen Lee, Owen Leung, Ellen Sai Nam Lo, Winnie Wing Yee Chan, Chun Kiu Lo, and veterinarians Jennifer L.L. Go, Hazel C.Y. Chung, and Kevin Cheng for taking care of lovebirds. We thank Cherry Pui Yu Wong, Chen Ma, Wynne Wan Hei Ting, Verna Wing Ting Shiu, Kit Ming Cai, Karson Cho Chun Fu, Katie Hei Yu Kwok, Cody Kwok Tsz Tseng, Isabella Lok Lum Wong, Bryan Chung Yin Leung for assisting the experiments. Special thanks to Xiaodong Wei for developing a video processing method using HPC, which greatly facilitated the efficiency of video combining and trimming. We thank Charis May Ngor Chan for technical support. This work was supported by a Start-up grant from the University of Hong Kong to S.Y.W.S.

## Supplementary Materials

### Supplementary figures

Figure S1-S4

### Supplementary video (available in the online version)

**Video S1.** A rosy-faced lovebird solving the combined puzzle, which includes the lid, the sliding door, and the drawer components.

---

**Figure S1.**
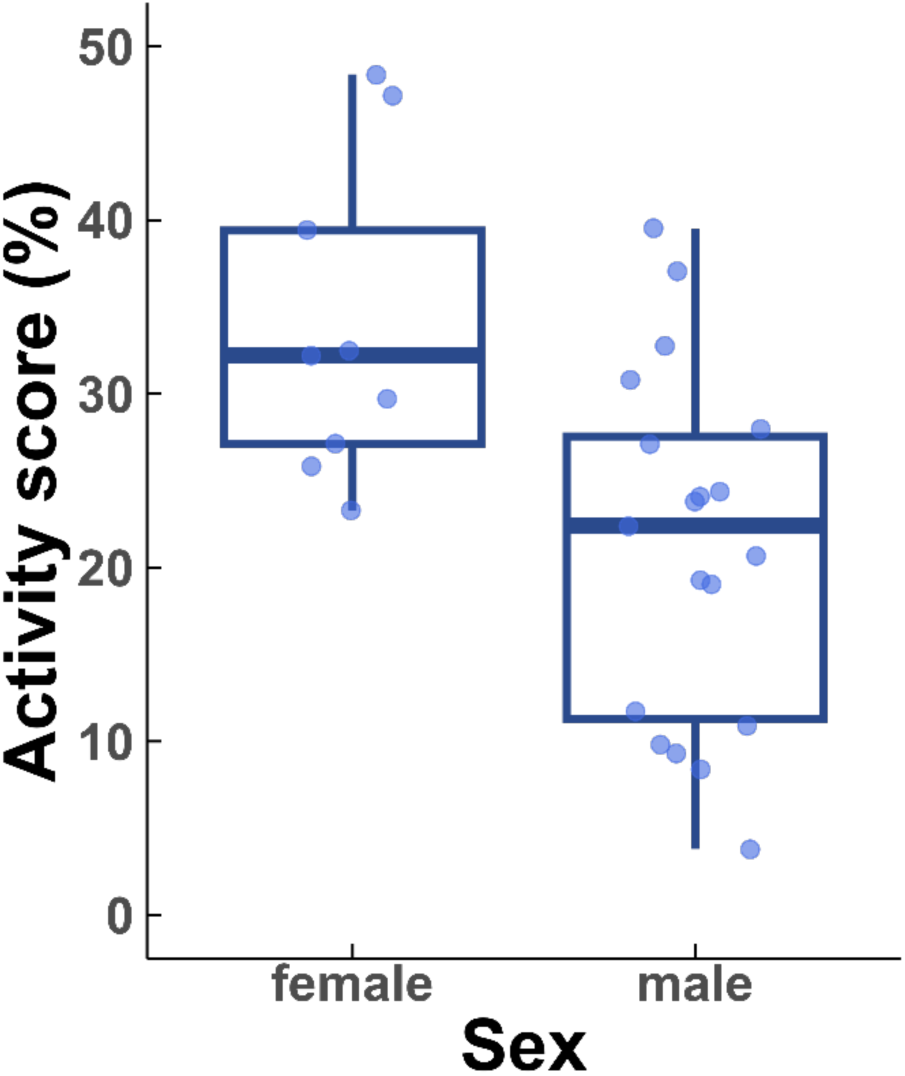
Sex differences in activity score.

**Figure S2.**
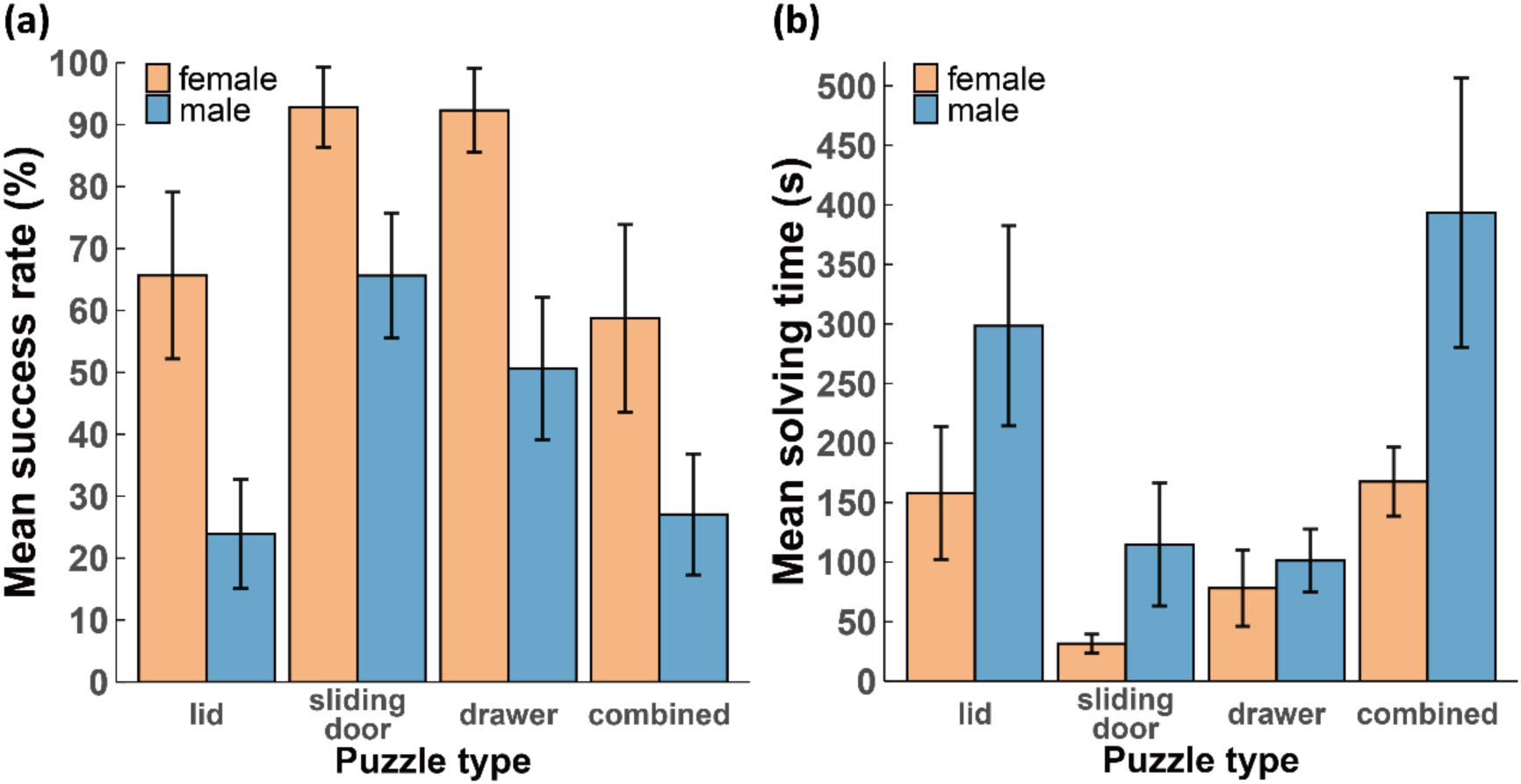
Sex differences in performance across four puzzles. (a) Mean success rate (±SEM) and (b) mean solving time (±SEM) by sex.

**Figure S3.**
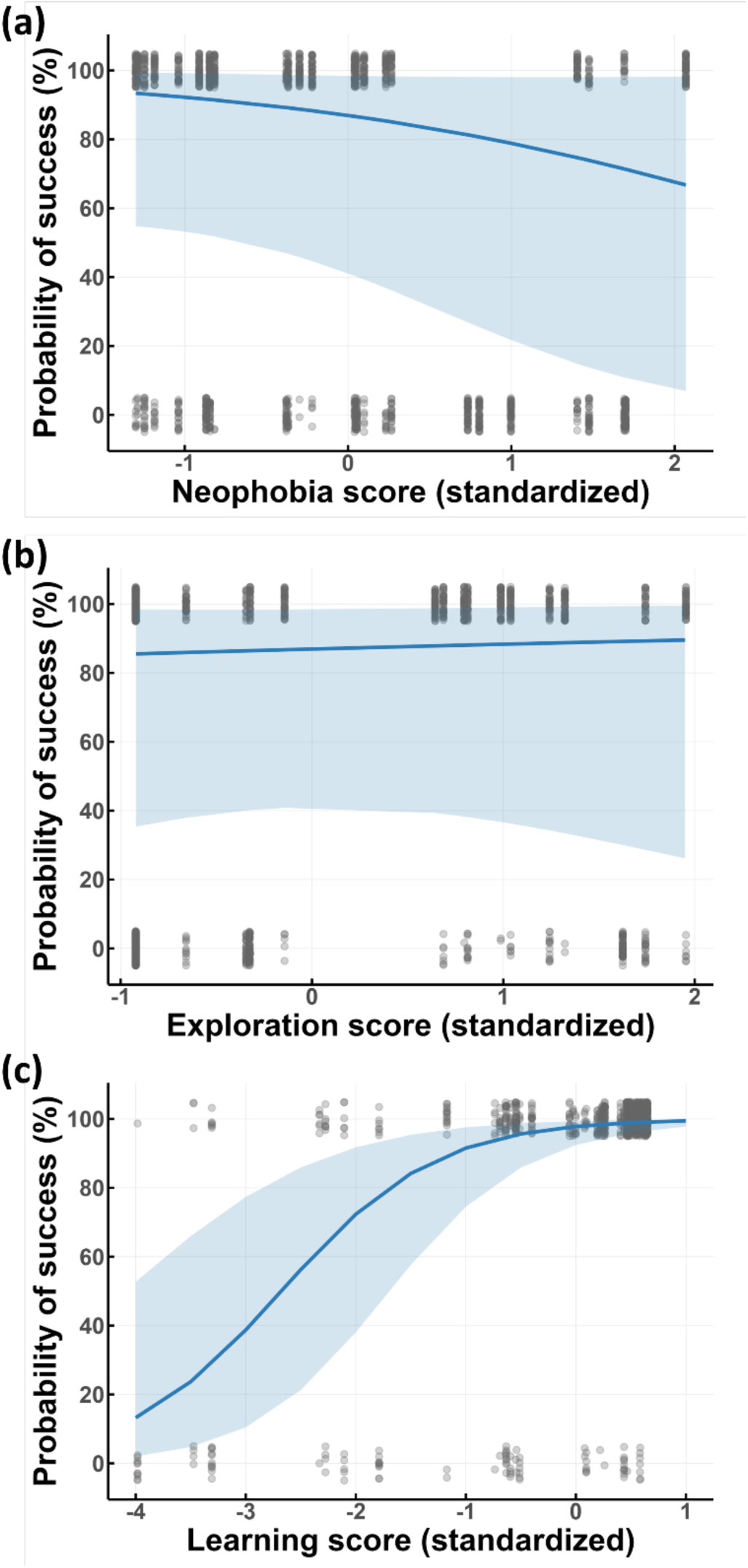
The association between puzzle-solving success and (a) neophobia score (Model 1), (b) exploration score (Model 1), and (c) learning score (Model 3).

**Figure S4.**
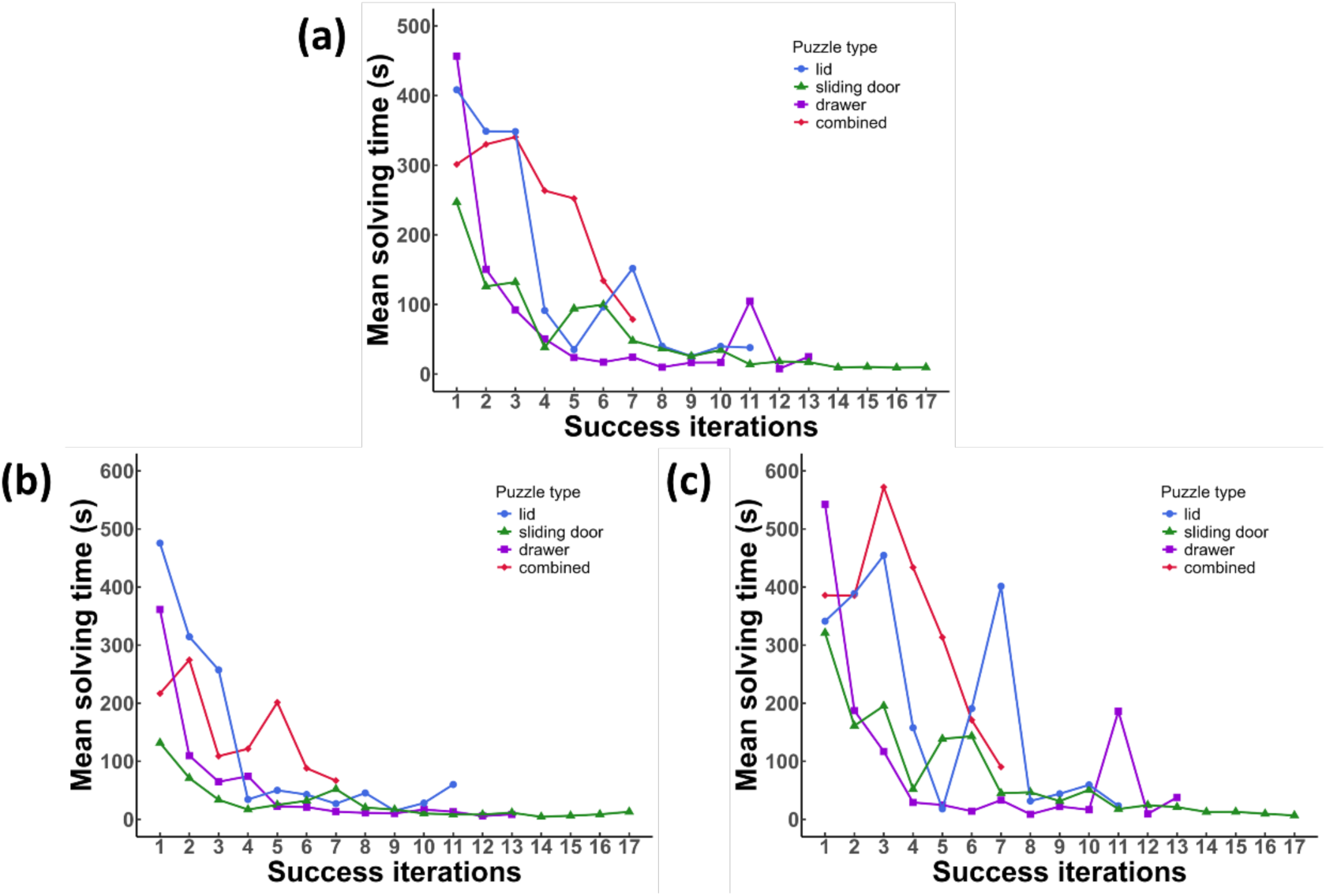
Learning curves across four puzzle types. Mean solving time over successive iterations for (a) all participants, (b) females, and (c) males.

